# PS Poly: A chain tracing algorithm to determine persistence length and categorize complex polymers by shape

**DOI:** 10.1101/2024.05.06.592664

**Authors:** Elizabeth A. Conley, Creighton M. Lisowski, Katherine G. Schaefer, Harrison C. Davison, Julie Baguio, Ioan Kosztin, Gavin M. King

**Affiliations:** Department of Physics and Astronomy, University of Missouri-Columbia, Columbia, MO 65211 USA; Department of Biochemistry, University of Missouri-Columbia, Columbia, MO 65211 USA; Materials Science and Engineering Institute, University of Missouri-Columbia, Columbia, MO 65211 USA

## Abstract

The fundamental molecules of life are polymers. Prominent examples include nucleic acids and proteins, both of which exhibit a large array of mechanical properties and three-dimensional shapes. The bending rigidity of individual polymers is quantified by the persistence length. The shape of a polymer, dictated by the topology of the polymer backbone, a line trace through the center of the polymer along the contour path, is also an important characteristic. Common biomolecular architectures include linear, cyclic (ring-like), and branched structures; combinations of these can also exist, as in complex polymer networks. Determination of persistence length and shape are largely informative to polymer function and stability in biological environments. Here we demonstrate <u>P</u>ersistence length <u>S</u>hape <u>Poly</u>mer (PS Poly), a near-fully automated algorithm designed to obtain polymer persistence length and shape from single molecule images obtained in physiologically relevant fluid conditions via atomic force microscopy. The algorithm, which involves image reduction via skeletonization followed by end point and branch point detection, is capable of rapidly analyzing thousands of polymers with subpixel precision. Algorithm outputs were verified by analysis of deoxyribonucleic acid, a very well characterized macromolecule. The method was further demonstrated by application to candidalysin, a recently discovered and complex virulence factor from *Candida albicans*. Candidalysin forms polymers of highly variable shape and contour length and represents the first peptide toxin identified in a human fungal pathogen. PS Poly is a robust and general algorithm. It can be used to extract fundamental information about polymer backbone stiffness, shape, and more generally, polymerization mechanisms.

## 1. INTRODUCTION

Knowledge of cellular function and dysfunction (disease) has advanced through developing a detailed understanding of many semi-flexible polymeric molecules. A prime example is the recently discovered peptide toxin candidalysin (CL), which is the virulence factor secreted by the fungus *C. albicans* ^1^. CL forms loops in solution which can then embed into membranes and form pores that damage host cells ^2^. When establishing the molecular basis of a disease, such as invasive candidiasis which emanates from *C. albicans* and has a high mortality rate ^3^, characterizing polymer mechanical properties and topology (shape) provides significant insight. For example, bending rigidity, quantified by the persistence length, sheds light on the expected diameter of loops formed by CL. This geometric information can be used to predict what size molecules might be able to pass through the CL pores in host cell membranes during *C. albicans* infection. Additionally, when studying the kinetics of polymer loop formation (cyclization) or branching, which usually involves secondary polymerization interfaces ^4,5^, it is informative to separate and quantify polymers by shape so that distinct reactions can be isolated. Such analyses can be used to construct kinetic models of looping and branching, to determine under what conditions polymer cyclization occurs, and to explore how conversion of linear polymers to looped polymers can be controlled.

Atomic Force Microscopy (AFM) is a powerful single molecule imaging technique employed in micro/nanoscale biophysical investigations and has been used to shed light on polymer persistence length, *l*_P_ ^6–9^. Analysis typically requires a stack of AFM image data containing many individual polymers. To perform *l*_P_ calculations, it is necessary to extract the coordinates along the chain contour, or “backbone”, for each polymer in the analysis. Once these coordinates are obtained, a polymer physics model such as the worm-like chain (WLC) can be used to deduce the persistence length through calculating mean-square end-to-end distances or correlations of backbone tangent vectors ^10^.

Existing software for polymer detection and characterization in AFM image data suffer from limitations ^11–21^. For example, popular tools including EasyWorm ^11^, Skan ^21^, and AutoSmarTrace ^19^, do not simultaneously provide robust feature extraction, skeletonization, shape, and mechanical property calculations such as persistence length. While EasyWorm is suitable for analysis of simple, unbranched polymer chains, it does not determine polymer shape, requires significant manual input, and is only available to the MATLAB community; thus limiting its accessibility and adaptability for open-source workflows. AutoSmartTrace, also MATLAB-dependent, requires manual intervention for thresholding and feature identification, reducing reproducibility across diverse polymer morphologies. While Skan provides Python-based skeletonization and network analysis, its focus on general image processing necessitates extensive customization for polymer-specific applications, lacking built-in modules for persistence length quantification. These limitations underscore the need for an open-source, flexible, modular, and easily extensible solution for the automated detection and analysis of polymer structures recorded through AFM imaging.

PS Poly, introduced here, attempts to fill these gaps and is well suited for studies of complex polymerization processes such as those underlying the host cell attack mechanism of *Candida albicans* ^2,22,23^. PS Poly is designed for polymer backbone isolation with sub-pixel precision, automated persistence length calculation, and shape categorization (e.g., linear, looped, branched, branched-looped, or overlapping chains). A workflow of the algorithm is shown (**Fig. 1**). The program is open-source with code written in both Python and Igor Pro 7 (WaveMetrics, Inc.) ^24^ and is near-fully automatic, requiring only basic information from the user such as the pixel resolution (nm/pixel) of the source images. PS Poly eliminates manual input post-thresholding, and outputs quantitative metrics such as total polymerized length and branch point coordinates. Persistence length results were verified by comparison to established values and were robust to moderate levels of added noise. The use of a Python framework may enhance accessibility, automation, open-source development, and modularity. Thus, the algorithm represents a step toward a standardized tool for AFM-based polymer analysis.

**FIGURE 1.**
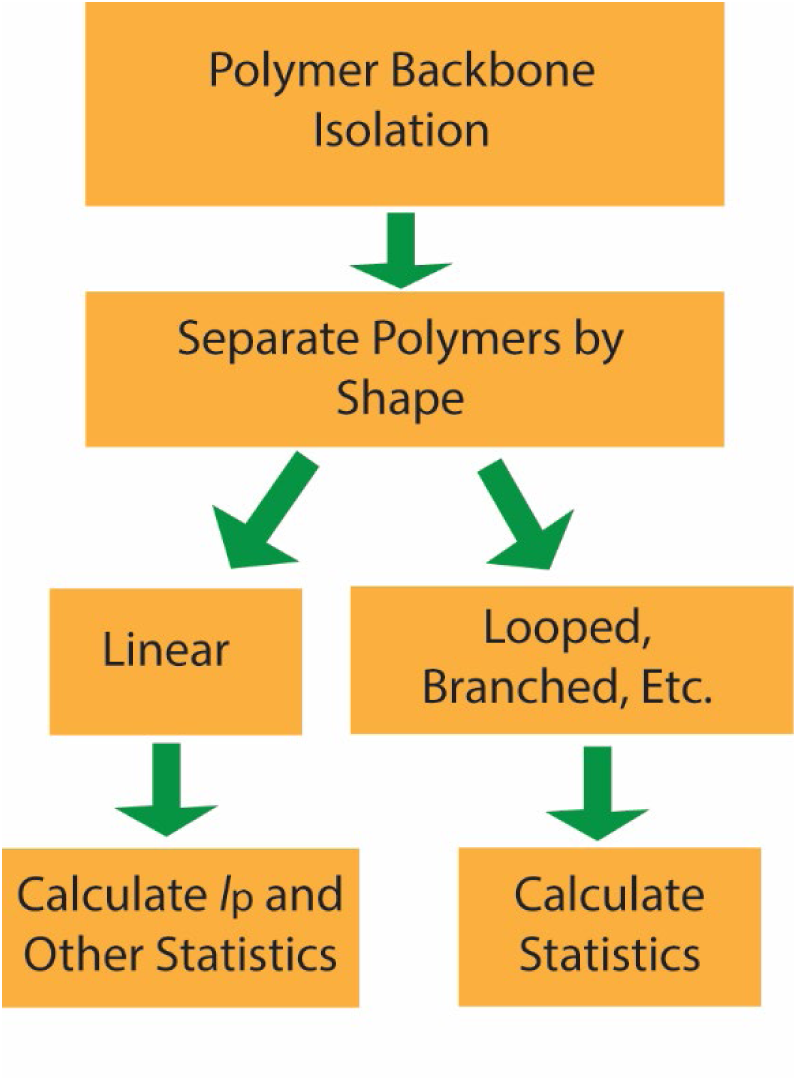
Program Overview. Images are processed and reduced to polymer backbone coordinates, then individual features are separated based upon shape. Usually, only measurements of linear polymers are considered when calculating persistence length, *l*_p_.

## 2. METHODS

In this section we first describe the algorithm as it was originally written in Igor Pro 7, subsequently referred to as PS-Poly. We then turn attention to the Python implementation, PsPolypy. The methods used for calculating the polymer persistence length and uncertainty are also discussed. Finally, we describe the techniques used for acquiring AFM images of CL in near-native conditions.

### 2.1 Igor Implementation: PS-Poly

Here we describe the Igor Pro 7 implementation, PS-Poly. Briefly, in this algorithm the images are skeletonized, a convolutional filter is used to identify endpoints, and a pathfinding algorithm is used to determine the coordinates of all linear particles used for persistence length analysis. To separate features by shape, filters were developed to identify branch points and distinguish branches from cyclic or looped polymers. These steps are described below. We assume that image preprocessing such as background subtraction has already been applied to the raw AFM image data prior to PS-Poly analysis.

#### Particle segmentation & polymer backbone isolation

The program begins by loading full-field images. Isolating the polymer backbone is the next step, illustrated in **Figure 2**. This is achieved automatically, by employing Otsu’s method ^25^, or manually, with user defined threshold corresponding to pixel intensity, which is proportional to topographical height, *z*, of the polymer in the AFM image. In either case, a binary “mask” image is created where all values above and below the threshold are set to 1 and 0, respectively. Then, if upscaling is desired, a copy of this mask is made with a higher pixel density by creating a new image with a specified scaling factor. The size of the expanded mask has dimensions scaled by the specified scaling factor, and each pixel in the original mask is taken up by a block of *x* scaling pixels in the expanded mask. This allows the result to be obtained with a subpixel level of accuracy which can be valuable for characterizing short polymers. A “skeleton” of the mask is created through a surface thinning algorithm which eliminates layers of the image until only single pixel linewidth traces remain (**Fig. 2D**) ^26^.

**FIGURE 2.**
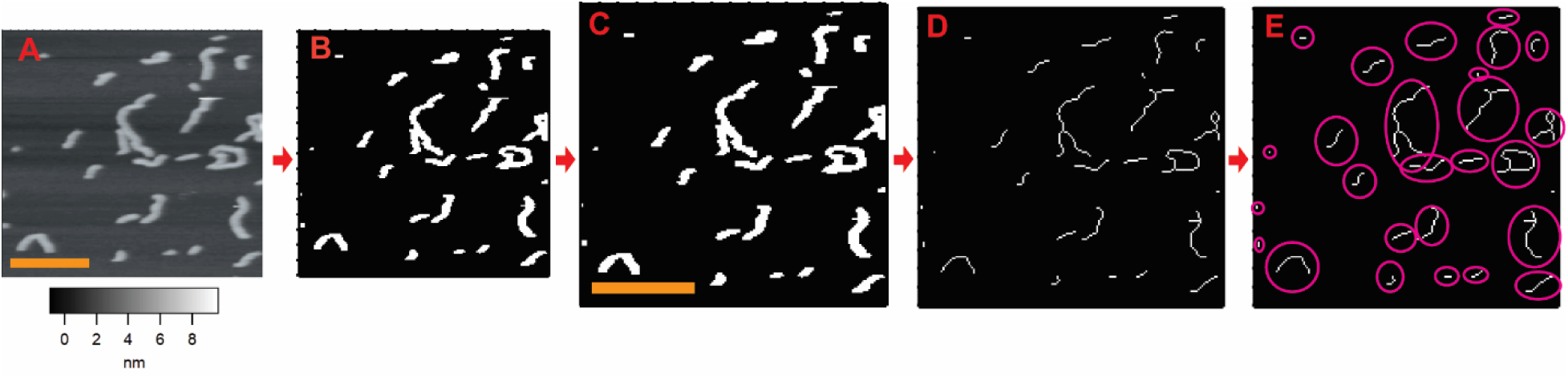
Polymer backbone isolation procedure. The first steps to perform PS-Poly calculations on an AFM image of candidalysin are shown. Scale bars (yellow) are 100nm. (**A**) Raw AFM image with topographic height, *z,* shown in greyscale. (**B**) Mask created from the AFM data. (**C**) New mask that has been expanded to a higher pixel density. (**D**) Skeleton created from the expanded mask. (**E**) Each molecule is registered as a unique object (pink circles).

#### Acquiring polymer coordinates

To obtain separate lists of coordinates for each molecule, we begin by looping through each pixel on a duplicate of the thinned image. Once a 1-valued pixel is found, that coordinate is stored.

Then a flood-fill algorithm fills-in all 1-valued pixels which are continuous with that coordinate. This process continues in a loop until the duplicated image is entirely 0-valued, and the resulting list of coordinates correspond to exactly one “seed” pixel per molecule. Then, depth-first search (DFS) is applied in a square around the seed pixel ^27^. DFS is used to identify connected pixels and provides detailed information about each connected component. The DFS algorithm explores all possible paths stemming from one input coordinate until a path is found to another input coordinate. The implementation of DFS in this program returns 1 if a path is found between the two coordinates, and 0 if there is no possible path. The search radius is incremented with each loop iteration, and the loop breaks once all locations continuous with the seed pixel are found. We found that applying DFS, as opposed to checking for continuity with all of the one-valued pixels in the image, reduces computational time significantly.

#### Sorting polymers by shape

Polymers are sorted based on their shape: Linear, Looped, Branched, and Branched-Looped. Examples of these four primary polymer classes are shown in AFM images of CL (**Fig. 3**). This sorting first requires polymer termination point identification, achieved through convolutional filtering.

**FIGURE 3.**
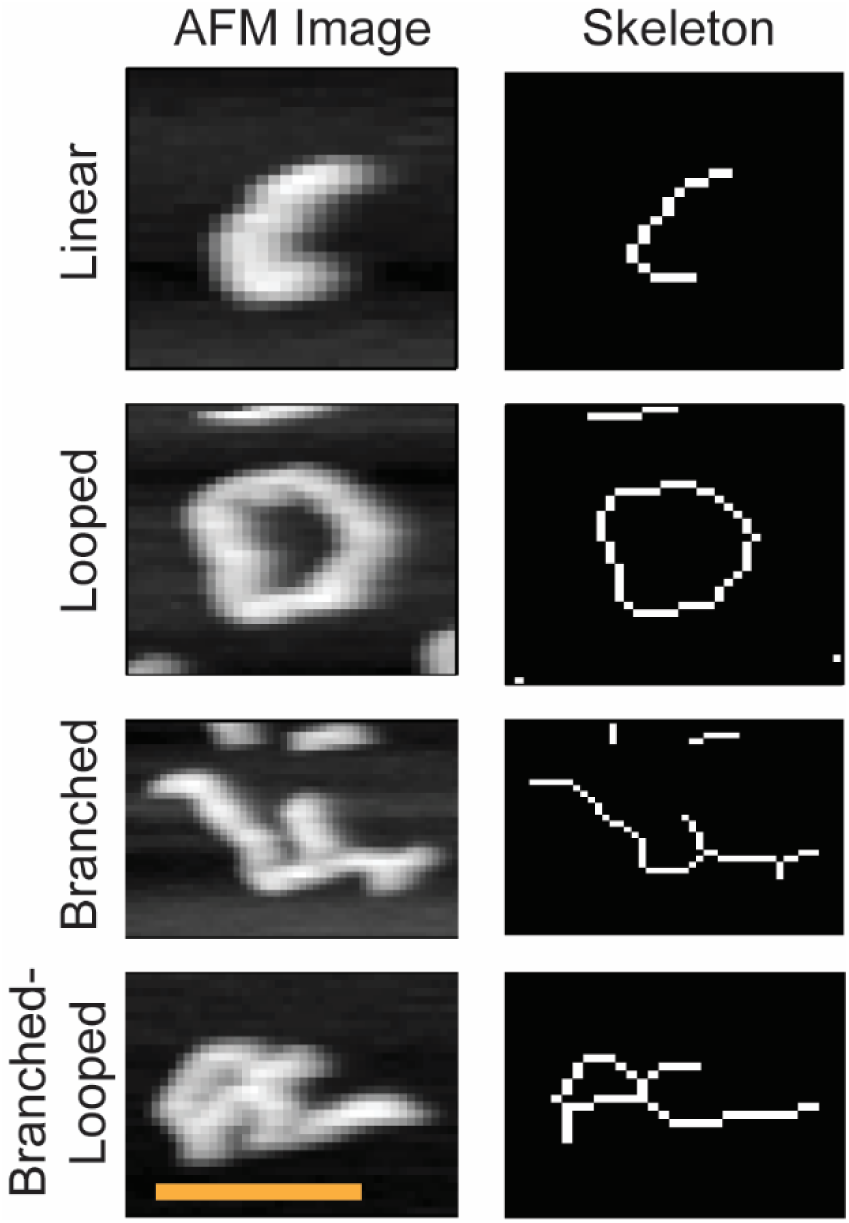
Primary polymer types. Examples of the different particle types identified through PS-Poly. Raw AFM image data of CL is shown next to processed skeleton images for Linear, Looped, Branched, and Branched-Looped polymers. The scale bar spans 50 nm and applies to all images.

#### Polymer end point determination

The algorithm loops through the image, cropping 9 × 9 pixel areas surrounding each central pixel test point. **Figure 4A** demonstrates the pixel grids that are created for every 1-valued pixel in the image. There are 16 possible endpoint configurations because there are 8 possible neighboring pixels, and so for one neighbor, there are 8 possible combinations. For 2 neighbors, there are 16 possible combinations, but only half of them are endpoints because the 2 neighboring pixels must also be adjacent to each other in order to be an endpoint. The pixel grids corresponding to the 16 possible endpoint configurations are shown in **Figure 4B**. Each pixel grid is compared with each of the 16 endpoint grids and if any one of them is an exact match, then that coordinate is considered to be an endpoint.

**FIGURE 4.**
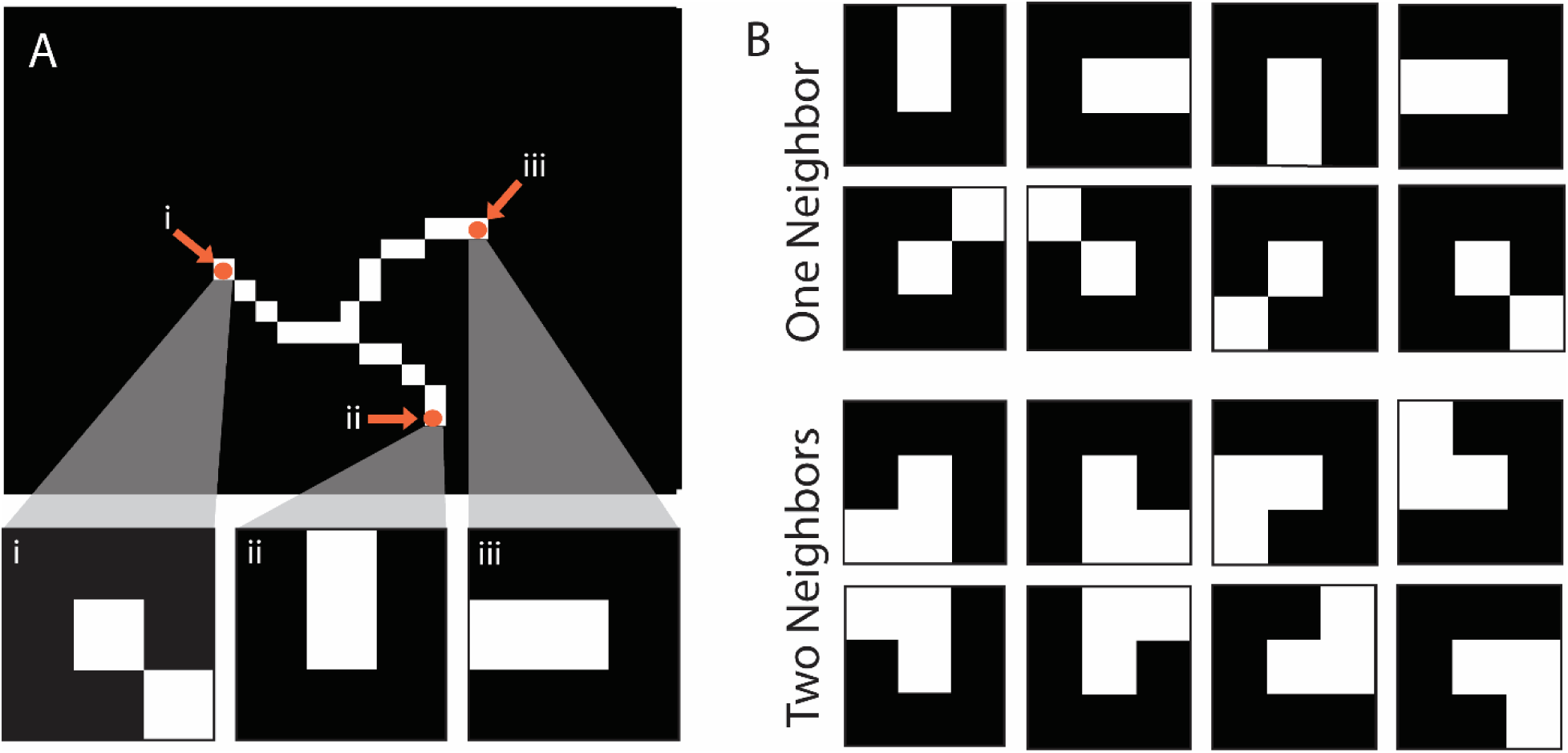
Convolutional filtering for end point determination. (**A**) shows 9 x 9 pixel grids that are created from points *i*, *ii*, and *iii* on a skeletonized polymer. (**B)** shows the sixteen different shapes that were used in the convolutional filter that finds endpoint coordinates. Shapes are clustered via number of neighbors.

#### Branch point identification

Following endpoint detection, branch points are identified through a filter that works by creating an array corresponding to neighboring pixel values that circle about a central pixel of interest (**Figure 5A**). Each path comprises a clockwise 10-pixel-long sweep. A unique path is counted for each 1 value that is abutted by 0 values (**Figure 5A**, blue triangles). We note that the last two pixels in the path are repeats of the first two, allowing evaluation of the starting point of the array. The array is examined for unique paths. If three or more unique paths are found, then the pixel of interest is determined to be a branch point. The algorithm is also prevented from overcounting branch points, as not all points with three neighbors are true branch points (**Figure 5B**).

**FIGURE 5.**
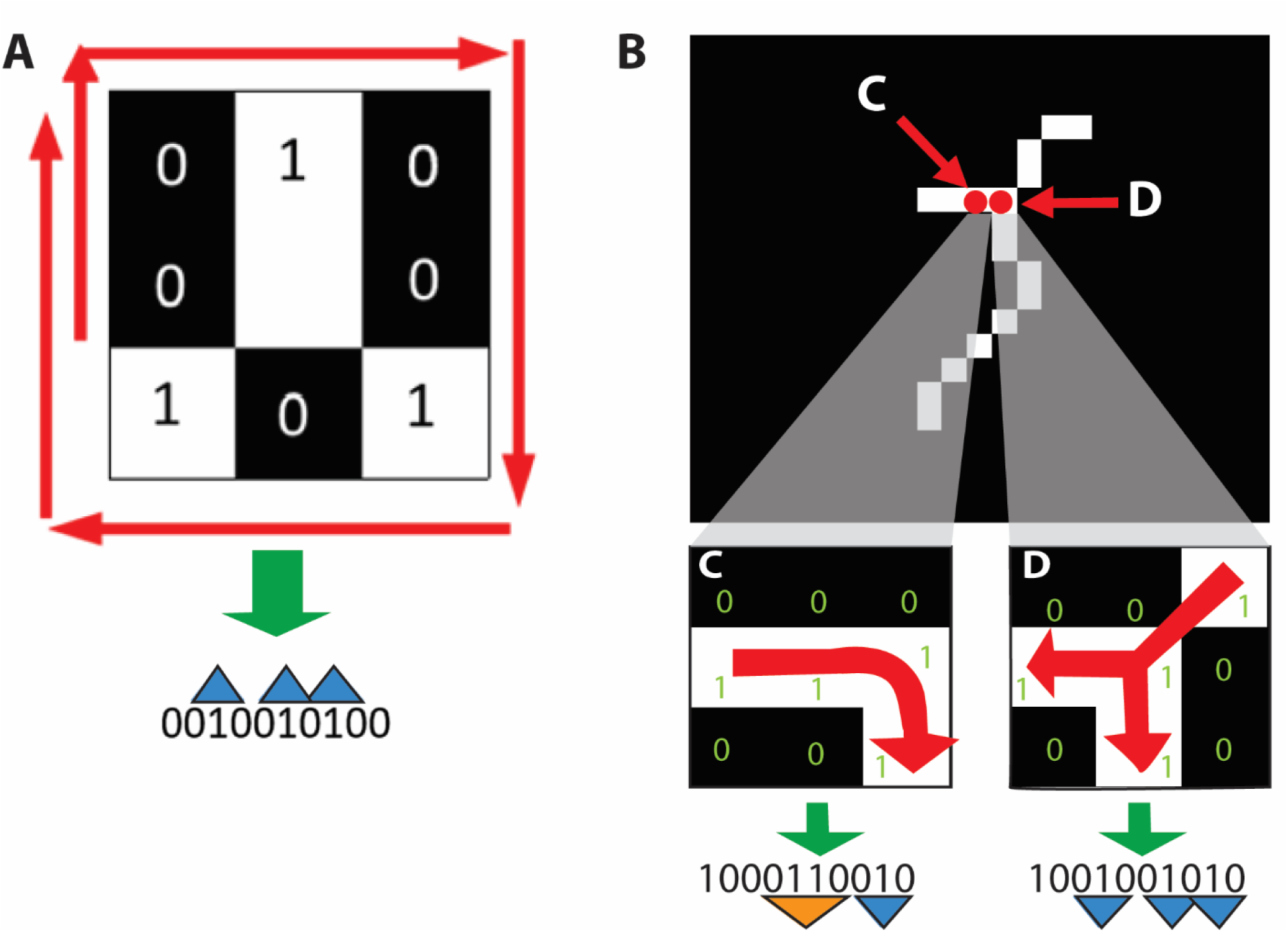
Branch point determination. (**A)** Each pixel in the image is centered in a 3 x 3 grid and a string of 1’s and 0’s begins at any point neighboring the test pixel. In this example three unique paths (blue triangles) are found, identifying the center pixel as a branch point. (**B)** Technique for prevention of branch point overcounting. Test point *D* has 3 neighboring 1-valued pixels, each of which corresponds to a unique path. Test point *C* also has 3 neighboring 1-valued pixels, but it does not exhibit three unique paths and is thus not a branch point. The non-unique path is marked by the orange triangle.

#### Shape calling, overlapped identification, and total polymer length determination

If a polymer has exactly two endpoints and no branch points, it is considered to be linear. If there are no endpoints and no branch points, then it is considered to be a loop. Further, polymers that branch and loop are separated from those which branch but do not loop. We further differentiated true branch points from segments where polymers drape over themselves as they adsorb to the imaging surface. These overlapped polymers were identified via co-localization of branch points with topographically high points along the polymer backbone (**Figure 6**).

**FIGURE 6.**
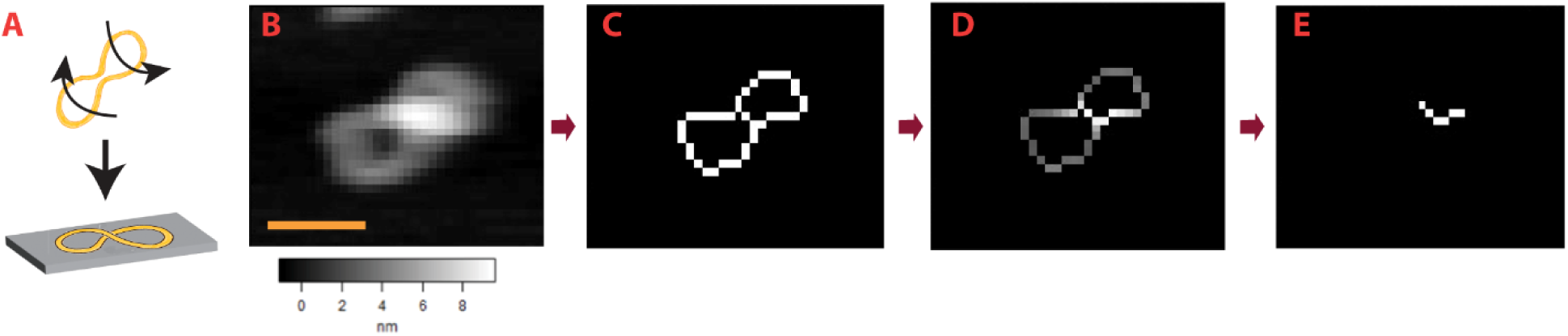
Process for determining overlapped polymers. (**A**) Cartoon showing an example of how an overlapped polymer may form. (**B**) AFM image of CL polymer. The height is shown in greyscale, the lateral scale bar (yellow) is 20 nm. (**C**) Skeleton created from the data in panel B shows a junction comprising the intersection of 4 branches. (**D**) New skeleton that incorporates height information. (**E**) Skeleton created at a threshold of 1.5 times the average height of the polymer distal from the junction identifies the polymer as overlapped.

The total polymerized length is found by applying a pathfinding algorithm which sums all pixels in each feature. For overlapped particles, the length is computed by adding the length of the original skeleton with the skeleton made at a threshold of 1.5 times the average height of all pixels on the skeleton with incorporated height information. If over 80% of the polymer is above 1.5 times the average height of the polymer backbone, then it is sorted separately as a noise particle. Such features could be aggregates or other artifacts. Polymers with high points that are not overlaps are still sorted by their shape and stored in a separate folder.

### 2.2 Python Implementation: PsPolypy

#### Preprocessing, Image Loading, & Upscaling

We assume that any image preprocessing including background subtraction has already been applied. PsPolypy uses as input a list of full-field images (denoted as *I*_*n*_) that contain polymer particles. The images *I*_*n*_ may have different pixel resolution (*N*_*n*_ × *M*_*n*_) but the same fixed real-space resolution (Res), measured in nanometers per pixel (nm/px). Each image is first converted to normalized grayscale, with pixel intensities 0 ≤ *I*_*n*_ ≤ 1. Optionally, users can upscale the pixel resolution of the images using *k*-order interpolation, as implemented in scikit-image ^28,29^. For a user-defined magnification factor *γ*, the pixel and the real-space resolution of the upscaled image becomes (*γ* · *N*_*n*_ × *γ* · *M*_*n*_) and Res/*γ* (nm/px), respectively. This optional step allows for finer image details to be analyzed.

#### Particle Segmentation

Each image *I*_*n*_ in the set undergoes particle segmentation through a multi-step process. Initially, Otsu thresholding is applied to create a binary mask *B*_*n*_. This mask then undergoes connected-component labeling, segmenting it into distinct regions, each corresponding to a unique polymer particle, with a well-defined bounding box. To ensure complete particle representation, any region whose bounding box touches the edge of the full-field image is discarded, as these particles may be partially cut off. The original image and binary mask are then cropped according to the bounding boxes of all remaining regions. Finally, a list of particle objects is created, with each object containing the cropped image of an individual particle and its corresponding cropped binary mask.

#### Skeletonization

The skeleton of each polymer particle (**Fig. 7**, *green*) is obtained by applying the skeletonize method from scikit-image to the corresponding binary mask. The resulting skeleton is then analyzed using Skan ^21^, which automatically determines the topology (e.g., liner, branched, looped or cyclic) and geometric features (e.g., end-to-end distance, contour length) of the particle. We denote the set of all paths as *P*, and the *m*-th path as *P*_*m*_. This skeletal analysis captures the main features of the particle’s structure and morphology.

**FIGURE 7.**
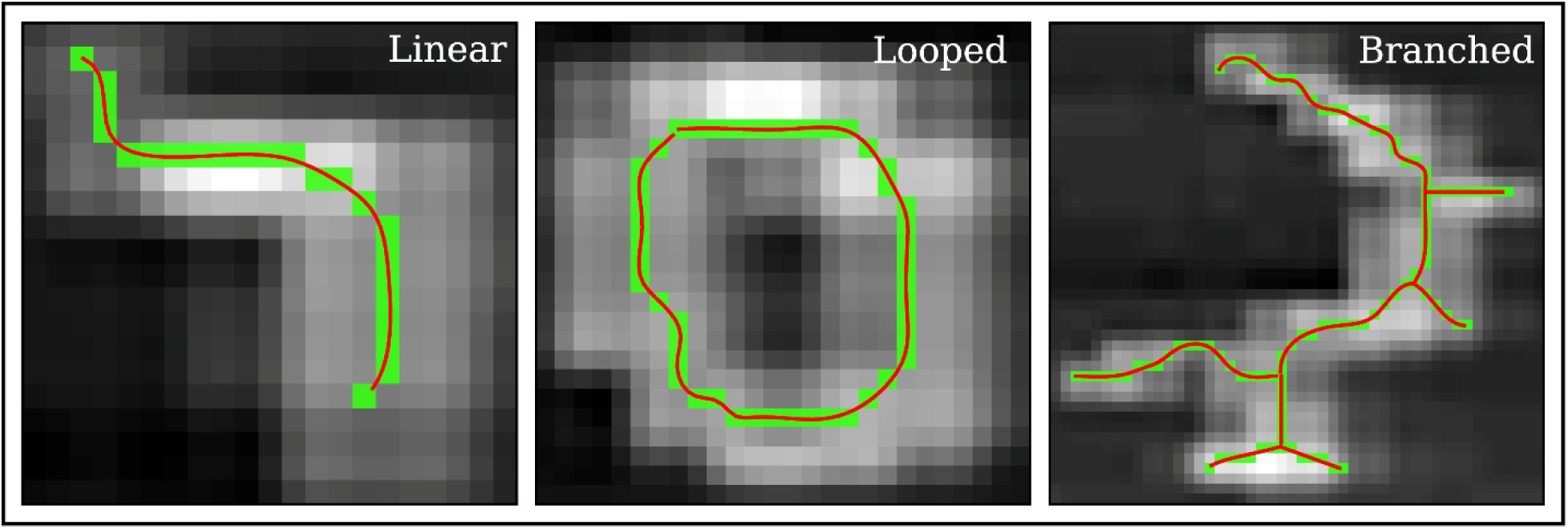
Skeletonization and interpolation of different polymer topologies. Examples of linear, looped, and branched polymers are shown along with the skeleton (*green*) and interpolation (*red*).

#### Classification

Polymers are classified into one of six categories based on their skeleton’s structure: Linear, Branched, Looped, Branched-Looped, Overlapped, or Unknown. If the skeleton contains a single path with distinct endpoints, the particle is classified as Linear (see **Fig. 7**). If the skeleton has a single path where the start and end points coincide, the particle is classified as Looped. For skeletons with multiple paths, each crossing is checked whether it is a branch junction or an overlap, defined by having a height 1.5 larger than the average polymer backbone height distal to the crossing. Polymers with at least one overlap are classified as Overlapped (as shown in **Fig. 6**). For the rest of the skeletons, if no combination of paths forms a cycle (as determined using the NetworkX graph representation of the skeleton) ^30^, the particle is classified as Branched. Conversely, if multiple paths are present and at least one cycle exists, the particle is classified as Branched-Looped. If none of the above criteria are met, the particle is categorized as Unknown. After classification, users may select only the types of particles they wish for further analysis.

#### Interpolate Skeletons

Each path (longer than 3 pixels) within each particle (digitized skeleton) undergoes cubic B-spline interpolation by employing the SciPy’s splev function ^31^. The coordinates along the interpolated skeleton path are given by *x*(*t*) and *y*(*t*), where *t* is the distance along the contour. The interpolated paths are sampled at user-defined intervals *dt* along the contour. The interpolated skeleton provides a more precise representation of the particle compared to the original (digitized) skeleton (**Fig. 7**, compare *red curve* with *green pixels*). For each sampling point along the interpolated path, both the position 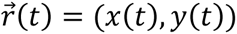 of the point and the tangent unit vector 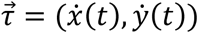 to the path are recorded.

#### Mean end-to-end distance

The mean end-to-end distance of the particles, ⟨*R*^2^⟩, as a function of contour length *L* is determined as follows. First, for the *m*-th path (*P*_*m*_), a symmetric distance matrix 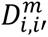 is constructed by calculating the Euclidian distance between all pairs of points *i* and *i*′ ≥ *i*, along the contour. Thus, the distances with the same lag *l* = *t*_*i*′_ − *t*_*i*_ correspond to the *l*-th superdiagonal of 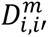. Finally, ⟨*R*^2^⟩ for *L* = *l* is calculated as the mean of the squares of all *l*-th superdiagonal elements of 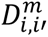 for all paths *P*_*m*_. The uncertainty of ⟨*R*^2^⟩(*L*) is estimated by calculating the standard error of the mean (SEM).

#### Mean orientation correlation function

The orientation (or tangent-tangent) correlation of polymer particles is defined by 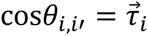 · 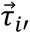, where 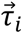 is the unit tangent vector to the interpolated path at point *i*. The mean orientation correlation ⟨cos*θ*⟩(*l*), as a function of path length *l*, is calculated by first constructing the correlation matrix 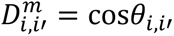, where *m* is the path index. For *l* = *t*_*i*′_ − *t*_*i*_, ⟨cos*θ*⟩(*l*) is calculated as the mean of all *l*-th superdiagonal elements of 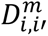 for all paths *P*_*m*_. Similarly to ⟨*R*^2^⟩, the uncertainty of ⟨cos*θ*⟩ is estimated through the corresponding SEM.

### 2.3 Persistence Length (l_p_) Calculations

In both Python and Igor Pro implementations, the persistence length, *l*_*p*_, of the polymer particles was estimated by fitting either ⟨*R*^2^⟩(*L*) or ⟨cos*θ*⟩(*l*) to their expressions from the worm-like-chain (WLC) model which is commonly used to describe semi-flexible polymers ^10^, i.e.,

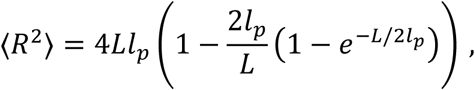

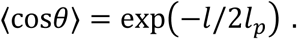

In PsPolypy, these nonlinear fits are performed using the feature rich lmfit python library ^32^.

### 2.4 Model Fitting and Uncertainty Analysis

In each data set {*x*_*i*_, *y*_*i*_, Δ*y*_*i*_}, where *x*_*i*_ represents the length along the paths, *y*_*i*_ the mean of either 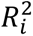 or cos*θ*_*i*_, and Δ*y*_*i*_ the corresponding standard error of the mean (SEM), we calculated the mean ⟨*y*⟩ and the uncertainty Δ*y* as a function of *x* = *l* or *x* = *L*. Data was fitted to the one parameter, *l*_*p*_, WLC model (discussed above) using the lmfit python package. The fitting process employed weighted least squares minimization, with weights calculated as 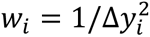. The best-fit parameter *l*_*p*0_ was determined by minimizing the chi-square statistic, with its uncertainty Δ*l*_*p*_ derived from the covariance matrix of the fit, scaled by the reduced chi-squared, 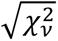. The 95% confidence interval (CI) for *l*_*p*_ was calculated as *l*_*p*0_ ± 1.96Δ*l*_*p*_, representing the range within which the true parameter value is likely to lie with 95% probability. To visualize the model’s predictive capability, 95% prediction bands were computed as f(x; l_p0_) ± 1.96 ∗ σ_total_(x), where f(x; l_p0_) is the WLC model of either *R*^2^or cos*θ* and 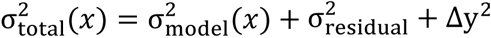. Here, *σ_model_* (*x*) represents the model uncertainty propagated from Δ*l*_*p*_, *σ_residual_* is the standard deviation of the residuals, and Δ*y* accounts for the measurement uncertainty. The resulting plots display the original data points with error bars ±Δ*y*_*i*_, the best-fit curve *f*(*x*; *l*_*p*0_), and the 95% prediction bands, illustrating both the uncertainty in the model and the expected range for new observations.

### 2.5 Atomic force microscopy imaging of CL

CL was purchased from Peptide 2.0 in powder, then hydrated to 100 μM in MilliQ water. The stock solution was stored in 10 μL aliquots at −80 °C. Imaging was performed as previously described ^2^. Briefly, a CL aliquot was thawed and diluted to 330 nM in the imaging buffer (10 mM Hepes, 150 mM NaCl, pH 7.3). Ninety microliters was added to freshly cleaved mica disks and incubated for 10 minutes at room temperature (∼25 °C). The samples were washed by exchanging 90 μL of imaging buffer five times to remove any particles in solution or loosely bound particles. Samples were imaged in the imaging buffer using biolever mini dips (Olympus, *k*∼0.1 N/m, *f_o_*∼30 kHz in fluid) in tapping mode (Cypher, Asylum Research). Throughout imaging, the tip sample force magnitude was kept to ≤100 pN, a regime in which minimal protein distortion is expected. Prior to algorithm implementation, images were flattened using commercial AFM software (Asylum Research).

## 3. RESULTS AND DISCUSSION

Double stranded DNA represents a convenient benchmark for polymer chain tracing algorithms as its persistence length has been well characterized. In an analysis of four AFM images containing 206 linear strands of DNA with data from Hennan *et al* ^8^, PS-Poly persistence length results for DNA were found to be 48 ± 3 nm. This is within the margin of error for the widely accepted value for double stranded DNA persistence length of around 50 nm (**Table 1**). The persistence length for the polymer Candidalysin was determined in an analogous manner and found to be 12.1 ± 0.3 nm using seven images containing 670 linear polymers. Using the Easyworm software ^11^, the results for Candidalysin were found to be 12 ± 3 nm^2^, which is in good agreement with our algorithm.

**Table 1.** Comparison of PS-Poly to previous work. Results were obtained via the end-to-end distance method for persistence length.

| Sample | Persistence Length,<br>Previous work | Persistence Length<br>using PS-Poly |
| --- | --- | --- |
| DNA | 46.8 +/- .6 nm<br>N = 126<br>(Ref 8) | 47.8 +/- 2.6 nm<br>N = 206 |
| Candidalysin | 12 +/- 3 nm<br>N = 191<br>(Ref 2) | 12.1 +/- .3 nm<br>N = 752 |

In the process of categorizing polymer feature shapes, the coordinates of all endpoints, branch points, and three-dimensional overlaps are stored in the output as well as the total length of each feature, total polymerized length for each image, and total polymerized length for all images. An example AFM image of CL and the resulting PS-Poly shape output are shown (**Fig. 8**).

**FIGURE 8.**
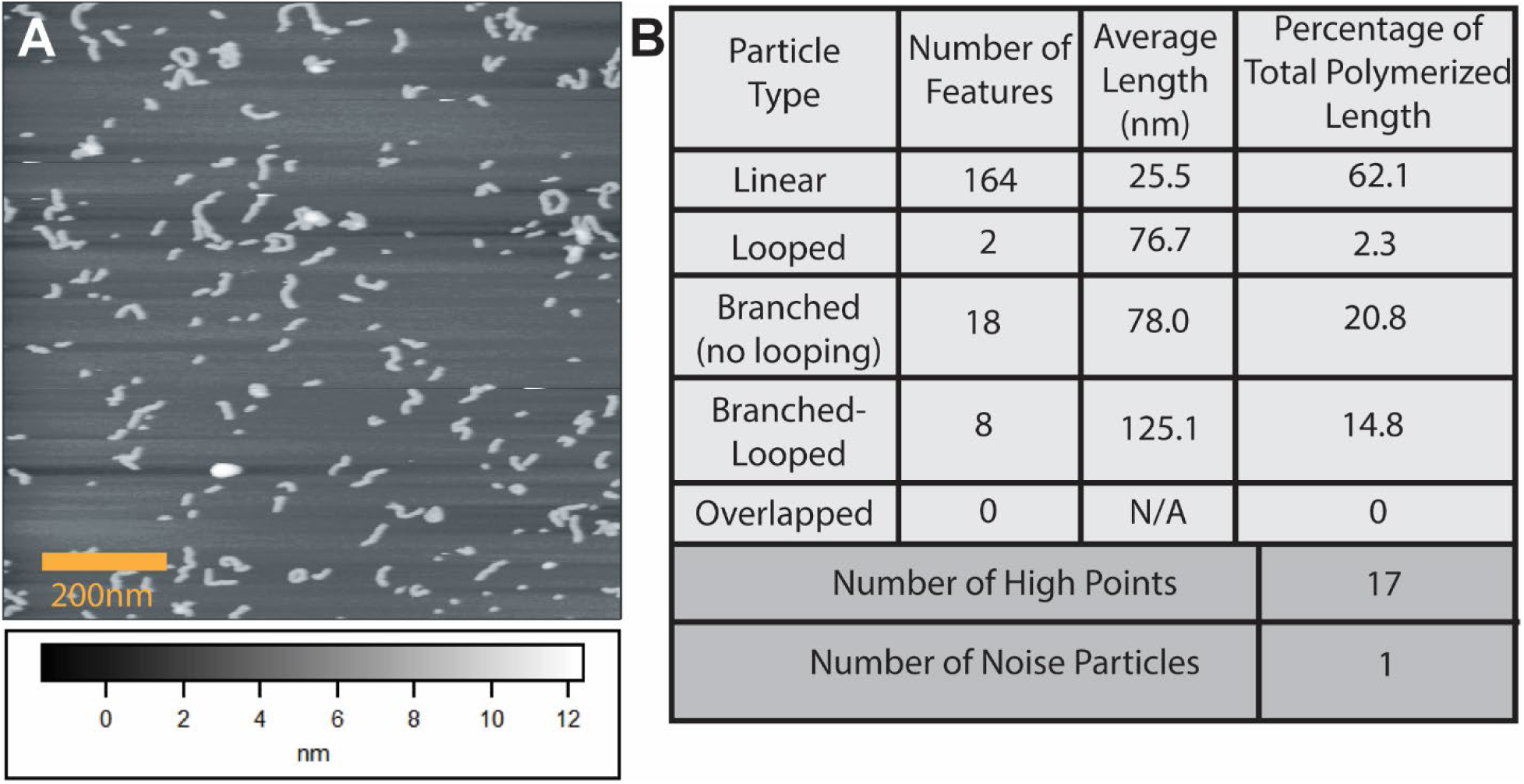
The output of PS-Poly for shape categorization. (**A)** Input AFM image of Candidalysin. The scale bar is 200 nm and the greyscale spans 12 nm. (**B**) Output for shape categorization shown in tabular format. Two types of artifact are identified, high points, defined as any point on a particle that is above 1.5 times the average height of the polymer backbone in the image and noise particles, defined as any particle in which 80% or more of the pixels are above 1.5 times the average height of the polymers in the image.

After establishing the overall agreement between our algorithm output and previous work, we analyzed the potential errors and robustness of the persistence length calculations. CL was employed as this polymer represents a general and complex test case. AFM image data revealed contour lengths ranging from individual CL subunits that appear as punctate features of dimension roughly equivalent to the AFM tip radius (∼5 nm) to well over 100 nm. However, CL polymers that are both Linear and long were rare. This is because the longer a CL polymer becomes the more likely it is to become Looped or Branched or both (i.e, Branched-Looped) ^22^. To handle these variations, in our CL *l*_p_ analysis we restricted the fitting window to contour lengths between 10 and 30 nm. The justification for excluding data with (a) *L* < 10nm, and (b) *L* > 30nm, is that the WLC model best describes semi-flexible polymers with *L* comparable or larger than *l*_*p*_, which in our case is larger than 10 nm; and (b) at higher contour lengths, the poor sampling of long CL polymers in the images makes the data analysis less robust. Only linear particles were used when fitting ⟨*R*^2^⟩ the WLC model. When considering only one end-to-end distance per particle, PsPolypy returned *l*_*p*_ = 12.0 ± 0.9 nm (**Fig. 9**). On the other hand, when considering multiple segments per particle, as described in the Methods section (which is the default), we obtained a more precise result, *l*_*p*_ = 12.5 ± 0.2 nm (**Fig. 10B**). In both cases, the fits were good quality, as indicated by *R*^2^ > 0.9 and the 95% prediction bands.

**Figure 9.**
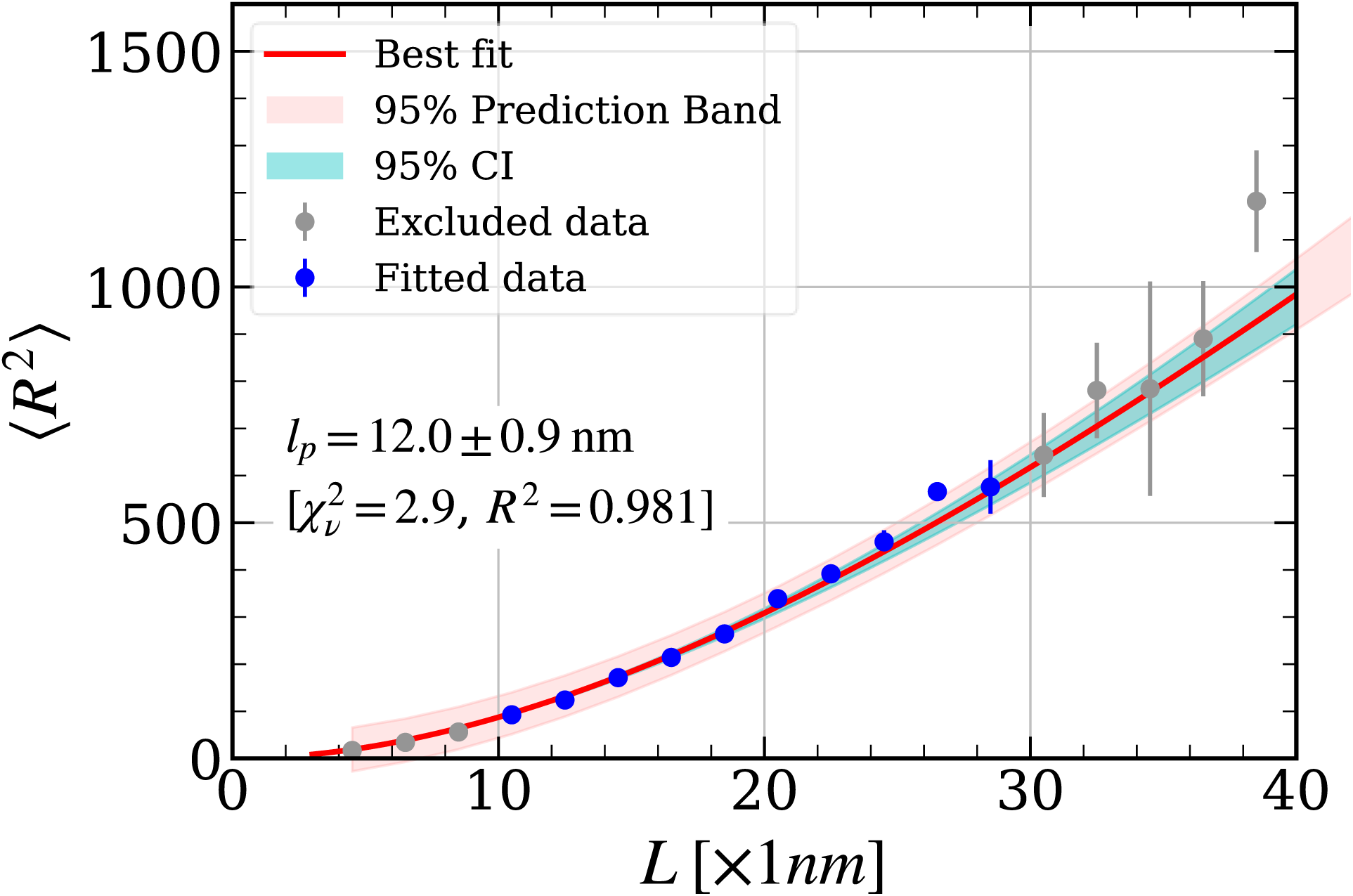
Limiting the contour length fitting window provides robust persistence length calculations for CL. Plot of mean-square end-to-end distance versus contour length for CL polymers. The persistence length *l*_*p*_ was found to be 12.0 ± 0.9 nm. A 95% confidence interval (CI, *teal shaded region*) and 95% prediction band (*pink shaded region*) are shown.

**Figure 10.**
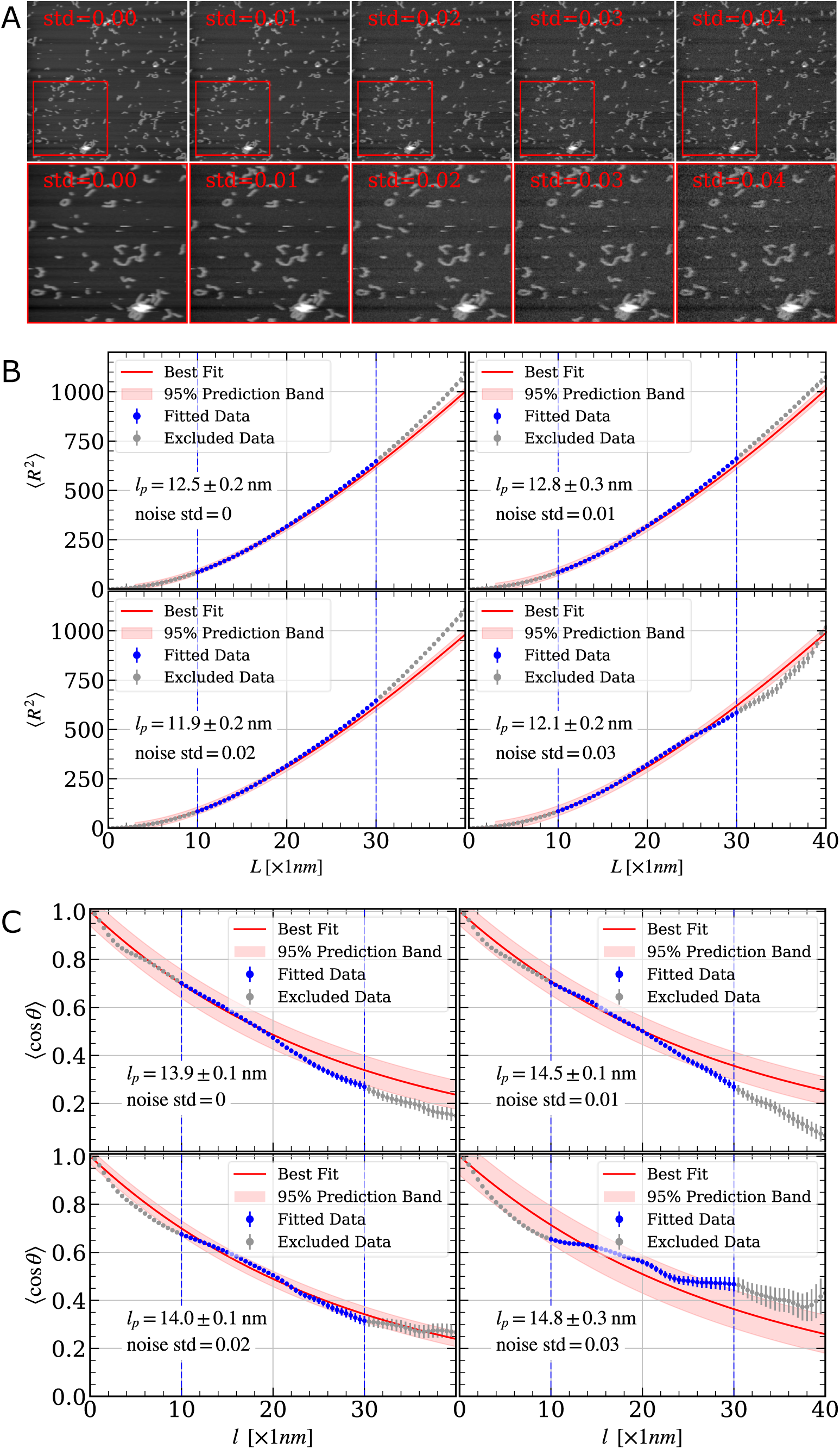
Persistence length calculations appear stable in the face of noise. (**A**) Image sequence showing the addition of noise. The second-row images are detailed views of the region indicated in upper images (*red rectangles*). The added noise levels for each plot are indicated and range between 0 and 0.03 (std). Calculations of persistence length using different methods: (**B**) end-to-end distance and (**C**) tangent-tangent correlation. As before, only data within 10 – 30 nm contour windows were included in the fits.

We next challenged the algorithm by repeating persistence length calculations with noise added to the raw image data. **Figure 10A** shows an AFM image of CL with an increasing amount of white gaussian noise added, quantified by the standard deviation (std). Analysis of the polymer persistence length from this data is shown using both the end-to-end distance (**R**^2^) method (**Fig. 10B**) and the tangent-tangent correlation (**TTC**) method (**Fig. 10C**). Despite this added noise, the calculated *l*_*p*_ value remains close to the nominal value of 12 nm (14 nm) using the **R**^2^ (**TTC**) method. Note that the slightly larger *l*_*p*_ value returned by the TTC method is most likely due to its sensitivity to the precise form of the interpolated skeletons. This conclusion is consistent with the relatively large 95% prediction band in **Fig. 10C**. It appears, however, that the algorithm is robust to a moderate amount of random pixel noise as is typically encountered in experimental settings.

The expected range for new observations are indicated by the 95% prediction bands. A summary plot displaying *l*_*p*_ outputs is provided in **Fig. 11**. Results from the **R**^2^ and **TTC** methods are shown in **Fig. 11A** and **B**, respectively. The tables contain additional information. We observe that as the image noise increases, the number of features detected as Linear decreases. This not only results in lower statistical weight of the *l*_*p*_ calculation but also in reduced quality of the fits.

**Figure 11.**
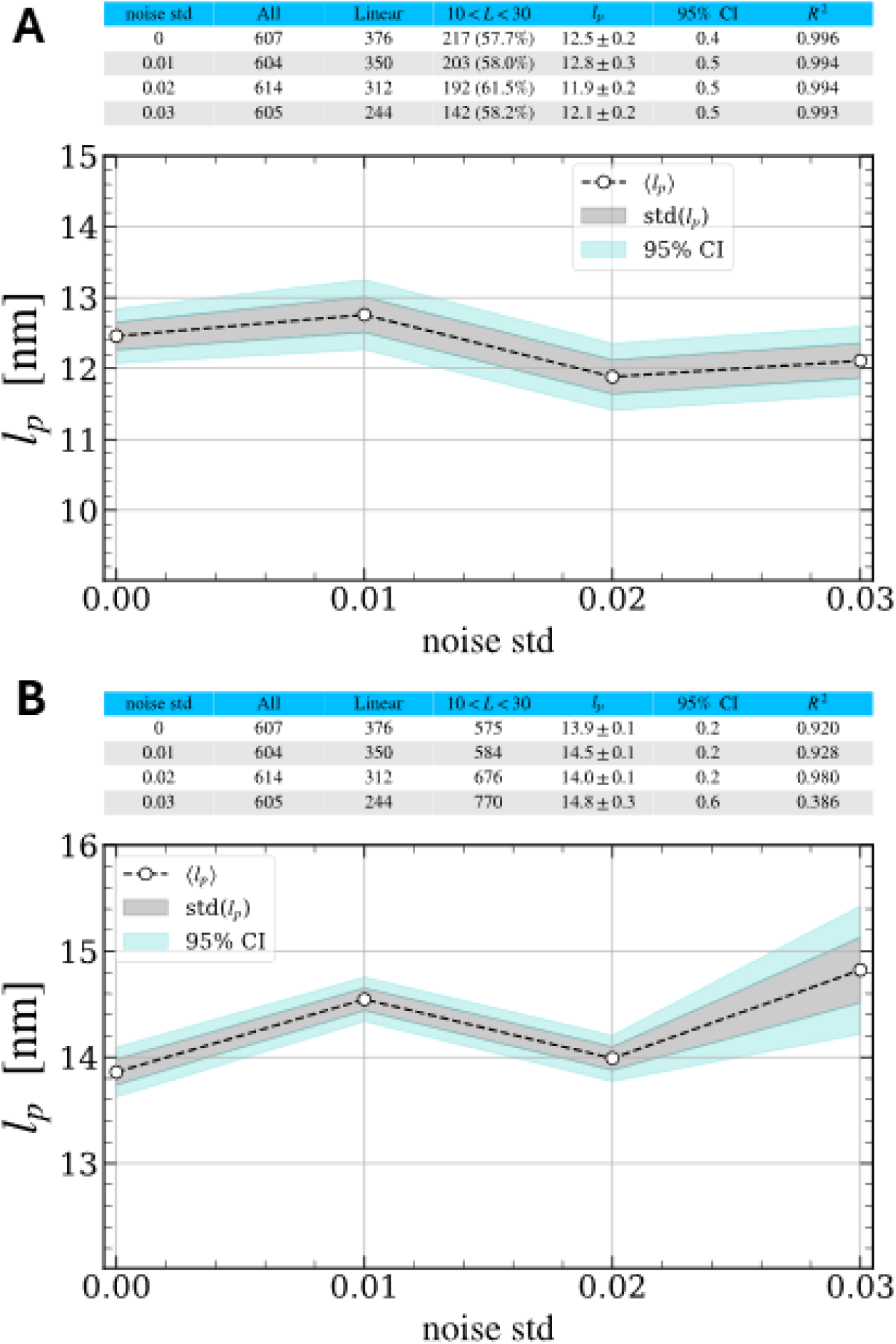
Persistence length output in the presence of noise. Persistence length results, as a function of added image noise, obtained by using the (**A**) **R**^2^ or (**B**) **TTC** methods. Rows in the tables are defined: ***noise std*** = noise level added to each image; ***All*** = total number of polymer features detected; ***Linear*** = number of features identified as Linear; ***10 < L < 30*** = number of Linear features exhibiting contour lengths within the 10 – 30 nm fitting window; ***l_p_*** = value of persistence length; ***95% CI*** = confidence interval; *R*^2^ statistical goodness of fit parameter.

### Computational Performance Evaluation

The runtime of PS-Poly in *Igor Pro 7* (64-bit) was compared to PsPolypy by processing a dataset of nine AFM images containing 779 particles using similar hardware. PS-Poly processed the dataset in 892 seconds on a CPU with an average frequency of approximately 4.2 GHz, while PsPolypy processed the dataset in 3.2 seconds on a CPU with an average frequency of approximately 4.4 GHz. While direct speedup comparisons are complicated by the variation of CPU utilization, these results suggest that the Python implementation achieves a significant reduction in runtime, on the order of two orders of magnitude. In addition to its superior calculation speed, Python’s open source and object-oriented design provides users with greater flexibility to fine-tune the algorithm to specific needs.

## 4. CONCLUSIONS AND FUTURE OUTLOOK

PS Poly is an algorithm designed to calculate both persistence length and shape from single molecule image data of complex polymers. When implemented as a python package the algorithm is modular and object oriented, making it straightforward to maintain and to extend its features and scope. While it can batch process stacks of AFM images, PsPolypy can also be easily used for customized workflows because most of the attributes and methods are directly available to the user. PS Poly currently provides two WLC model-based methods for calculating persistence length (**R**^2^ and **TTC**) from the skeletonized representation of the polymer particles; additional methods can be added in the future, as needed. Furthermore, its automated shape detection can be used to provide insight into polymerization mechanisms, as we have recently shown ^22^. The benefits of automating this process include reduced human bias as well as time saved by the user and the related improvement in statistical weight of the data set. The current implementations of PS Poly use the Otsu’s method for image thresholding which works effectively for high signal-to-noise images such as those typically acquired in AFM. However, other segmentation methods such as watershed algorithms and machine/deep learning features can also be added to future versions. For best results, it is important to reduce noise in the input images before running the program. As we showed, the algorithm is robust to moderate levels of random pixel noise, but it will break down if higher noise levels are encountered. While PS Poly was developed to analyze AFM image data, it has the potential to generalize to other imaging modalities.

